# Scene segmentation processes drive EEG-DCNN alignment

**DOI:** 10.1101/2025.03.11.642570

**Authors:** Jessica Loke, Amber M. Brands, Natalie Cappaert, H. Steven Scholte

**Affiliations:** Department of Psychology, University of Amsterdam, Amsterdam, The Netherlands; Amsterdam Brain & Cognition (ABC) Center, University of Amsterdam, Amsterdam, The Netherlands; Video and Image Sense Lab, Informatics Institute, University of Amsterdam, Amsterdam, The Netherlands; Swammerdam Institute for Life Sciences, University of Amsterdam, Amsterdam, The Netherlands

## Abstract

Visual processing in biological and artificial neural networks has been extensively studied through the lens of object recognition. While deep convolutional neural networks (DCNNs) have demonstrated hierarchical feature extraction similar to biological systems (DiCarlo and Cox, 2007; Yamins and DiCarlo, 2016), recent findings reveal a growing discrepancy: DCNNs with higher object categorization accuracy paradoxically show worse performance at predicting neural responses (Xu and Vaziri-Pashkam, 2021; Linsley et al., 2023). Using a large-scale human electroencephalography (EEG) dataset (*n*=10, 82,160 trials), we investigate whether this discrepancy arises because human neural EEG signals predominantly reflect scene segmentation processes rather than high-level, category-specific object representations. We trained DCNNs to perform object recognition using visual diets (∼1 million training images across 292 object categories) with systematically varying scene segmentation demands: objects-only (pre-segmented), background-silhouette (explicit boundaries), original/background-only images (requiring full segmentation). Despite substantial differences in categorization accuracy (27-53%), all trained models showed remarkably uniform encoding performance, with peak correlations with neural data at ∼0.1s post-stimulus. Layer-wise analysis revealed a significant negative correlation between categorization accuracy and encoding performance, with earlier network layers better predicting EEG responses than deeper layers specialized for object categorization. This dissociation suggests that EEG signals primarily reflect fundamental scene parsing mechanisms rather than object-specific representations, explaining the growing discrepancy between DCNN’s increasing categorization performance but deteriorating neural prediction performance.

**Significance Statement:** This research provides a novel perspective on human electroencephalography (EEG) signals during visual processing through systematic manipulation of scene segmentation demands in deep neural networks. Using a large-scale dataset of 82,160 EEG trials and 20 trained DCNNs, we demonstrate that EEG responses primarily reflect early visual processing involved in breaking down and organizing visual scenes (scene segmentation/parsing) rather than high-level object recognition. This finding helps explain previously observed discrepancies between DCNNs’ categorization performance and neural prediction accuracy, suggesting that improving models’ ability to segment scenes, rather than simply recognizing isolated objects, may better align artificial and biological visual processing.

## Introduction

Visual processing in both biological systems and artificial neural networks, such as deep convolutional neural networks (DCNNs), is associated with the ability to categorize objects from visual inputs (Kriegeskorte, 2015). Early comparative studies showed that DCNNs which are optimized for object categorization tasks extract visual features hierarchically, in a manner similar to biological neural systems (DiCarlo and Cox, 2007; Khaligh-Razavi and Kriegeskorte, 2014; Yamins et al., 2014; Yamins and DiCarlo, 2016). These models replicated visual processing hierarchies observed in humans, both in anatomical region-based processing steps (Güçlü and van Gerven, 2015) and temporal dynamics in EEG responses (Yamins et al., 2014; Cichy et al., 2016). Crucially, these findings further demonstrated that DCNNs’ performance on object categorization aligns closely with DCNNs’ ability to predict neural data. Consequently, DCNNs that reach categorization accuracies closer to human-level performance are expected to more accurately predict human (and also non-human primates) neural data. However, recent studies have identified a puzzling disconnect: improvements in DCNNs’ object categorization performance no longer translate to enhanced neural prediction accuracy (Xu and Vaziri-Pashkam, 2021; Linsley et al., 2023). As DCNNs achieve increasingly higher performance on benchmarks like ImageNet, their ability to predict human neural responses has plateaued or even declined. This suggests that modern DCNNs may be optimizing features that diverge from those used by biological visual systems.

This disconnect points to a fundamental question: what aspects of visual processing might DCNNs be missing? One crucial difference between biological vision and current DCNNs lies in how they approach scene understanding. While DCNNs excel at categorizing pre-segmented objects, biological vision must first parse complex scenes into meaningful components before object recognition can occur (Roelfsema, 2006). This process of scene segmentation—which involves detecting boundaries, separating figures from backgrounds, and integrating local features with global context—poses unique computational challenges that aren’t well-captured by feedforward networks optimized for ImageNet-style tasks (Kietzmann et al., 2019). Unlike object recognition, which can potentially be accomplished through feedforward processing alone, scene segmentation requires more complex recurrent processing to resolve ambiguities and integrate contextual information (Lamme and Roelfsema, 2000; Scholte et al., 2008; Pitts et al., 2012; Self et al., 2013; Seijdel et al., 2020; Loke et al., 2022). Standard datasets like ImageNet have predominantly featured objects against uniform backgrounds, effectively presenting “pre-segmented” objects. This simplification may have led to an overemphasis on high-level object features while underestimating the importance of scene segmentation processes in visual processing. Recent work suggests that early and intermediate stages of visual processing, which extract features such as edges, textures, and shapes, may play a more fundamental role in visual perception than previously thought (Zheng et al., 2019; Loke et al., 2024).

In this study, we explore whether the discrepancy between DCNNs’ performance and neural prediction might be explained by scene segmentation processes. We systematically vary scene segmentation demands across different training conditions to investigate how they affect both object categorization performance and alignment with neural responses. Using a large-scale electroencephalography EEG dataset (n=10, 82,160 trials), we test whether neural signals predominantly reflect scene segmentation processes rather than high-level object representations. By training models on four distinct visual diets—original images (requiring full segmentation), object-only images (pre-segmented), background-silhouette images (explicit boundaries), and background-only images—we can dissociate the contributions of scene segmentation from object recognition and examine their relative importance in predicting neural responses. This systematic approach allows us to directly test whether the ability to predict neural responses relies more on capturing scene segmentation processes than on object recognition performance. We hypothesize that models trained with higher segmentation demands will show enhanced alignment with neural responses, even if they achieve lower categorization accuracy.

## Materials and methods

### Image Datasets

We developed four distinct image conditions (see Figure 1) from the Google Open Images V5 Dataset (Krasin et al., 2017; Benenson et al., 2019) to systematically investigate the role of scene segmentation in visual processing. The original condition contained unaltered images with objects in their natural contexts. For the object-only condition, we isolated target objects against a black background using the dataset provided segmentation masks. The background-silhouette condition preserved the original backgrounds with object silhouettes, while the background-only condition removed object silhouettes using the LaMa inpainting algorithm (Suvorov et al., 2021). Each condition comprised 292 object classes, with 941,235 training images and 53,239 test images. We excluded object classes with fewer than 100 training and test images to ensure robust training. For images with multiple objects, we designated the class of the largest object as primary.

**Figure 1.**
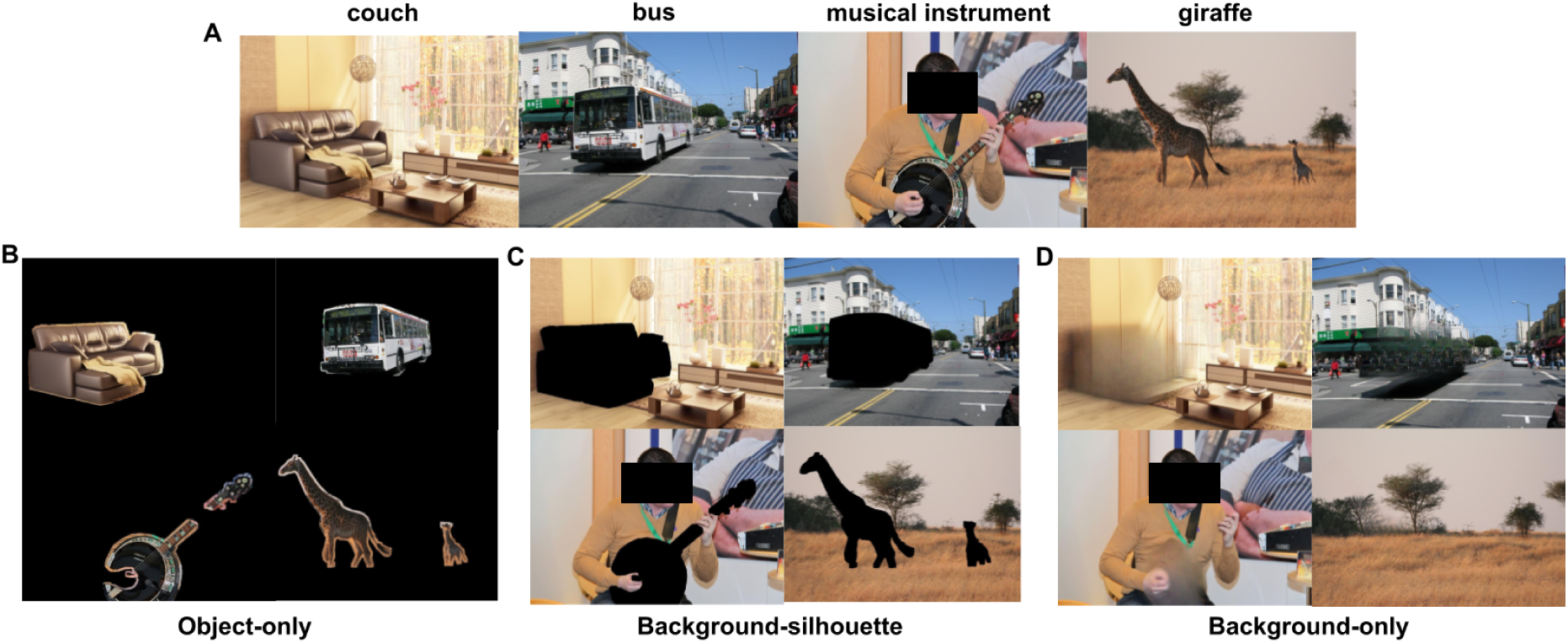
Visual diet conditions for DCNNs training. A) Original images from Google Open Images V5 Dataset showing objects in their natural backgrounds. B) Object-only condition displaying target objects isolated against a black background, created using dataset segmentation masks. C) Background-silhouette condition showing object silhouettes presented against their original backgrounds. D) Background-only condition with target objects removed and the backgrounds preserved through inpainting.

For fine-tuning and electroencephalography (EEG) prediction, we employed the THINGS Image Dataset (Hebart et al., 2019) which contains naturalistic object images across 1,854 common object concepts. This dataset was chosen because it was used during the EEG recordings made by (Gifford et al., 2022) (see section below). The image subset used for the EEG recordings contained 1,854 object classes, with 16,540 training images and 200 test images.

### EEG Dataset and Preprocessing

We utilized a large-scale EEG Dataset (Gifford et al., 2022) recorded from 10 human participants. Each participant completed 82,160 trials. The experimental paradigm included a training set of 16,540 images (each presented 4 times) and a test set of 200 images (each presented 80 times). Participants performed an orthogonal target detection task while viewing the images. The data were preprocessed following the original study’s preprocessing pipeline, retaining only 17 occipital electrodes (for emphasis on visual processing) with trials epoched between -0.2 and 0.8s.

### Deep convolutional neural networks architecture and training

We trained 20 deep convolutional neural networks using the AlexNet architecture (Krizhevsky et al., 2012), implementing five initializations for each of our four image conditions. We chose AlexNet for its established performance in predicting EEG responses and computational efficiency. Multiple initializations were used to assess variance in its internal representations (Mehrer et al., 2020) The models were trained using a cross-entropy loss function with stochastic gradient descent (momentum = 0.9) and a learning rate of 0.005, which decayed by 0.1 every 30 epochs. Training continued for 80 epochs with a batch size of 256. For comparison purposes, we included five randomly initialized AlexNet models and five AlexNet pre-trained on ImageNet as controls.

### Feature extraction and encoding models

We extracted model activations from eight layers spanning early to late processing stages: maxpool1, maxpool2, ReLU3, ReLU4, maxpool5, ReLU6, ReLU7, fc8. Given the inherently nonlinear transformations occurring across layers, we opted for a KernelPCA because it allows us to capture the complex, nonlinear relationships in unit activations across layers. We applied KernelPCA with a 4th degree polynomial kernel for nonlinear dimensionality reduction on the extracted features. A total of 1,000 principal components were obtained per image, based on a computation and model optimization tradeoff.

We constructed linearized encoding models to map the extracted features to EEG responses for each channel and time sample (Holdgraf et al., 2017; Gifford et al., 2022). The stimulus features were represented by a predictor matrix of 16,540 (number of images) x 1,000 (principal components). We averaged EEG responses across the four repetitions of each training image, then regress the EEG amplitudes (per channel and time sample) onto our predictor matrix using a ridge regression with a built-in cross-validation (sklearn.linear_model.RidgeCV). The alpha parameter for ridge regression was optimized using leave-one-out cross validation across seven values (np.logspace(-1,5,7)). Subsequently, the regression weights obtained from the training images are then used to predict the EEG amplitudes for 200 held-out test images. The encoding models are fitted per participant - i.e. regression weights are trained and tested within the same participant.

### Statistical analysis

#### Comparison of predicted EEG with recorded EEG responses

To evaluate encoding performance, we randomly split the 80 repetitions of recorded EEG responses to test images in half and computed Pearson correlations (scipy.stats.pearsonr) between the mean of 40 randomly chosen repetitions and the predicted EEG amplitudes. This procedure was repeated 50 times. The averaged correlation values across 50 iterations and across all EEG channels were reported.

#### Determination of noise ceiling

For the upper bound of the noise ceiling, we randomly split the 80 repetitions of recorded EEG responses to test images in half and correlated the mean of 40 randomly chosen repetitions with the mean of all 80 repetitions. For the lower bound of the noise ceiling, we correlated the mean of 40 randomly chosen repetitions with the mean of the other 40 repetitions. Similar to the evaluation of the predicted EEG responses, we repeated the noise ceiling computation 50 times. The averaged correlation values across 50 iterations determined the noise ceiling bounds.

### Statistical tests

#### Model categorization performance comparison

To evaluate differences in final validation accuracy among trained models, we performed pairwise t-tests (scipy.stats.ttest_ind). The statistical significance of these comparisons was established using an alpha level of 0.05 adjusted by Bonferonni correction.

#### Model encoding performance comparison

To evaluate models’ encoding performance, we derived two metrics: the peak correlation between models’ predicted EEG amplitudes and the actual EEG recordings, and the area under the curve (AUC) of these correlation values across all time samples. Pairwise t-tests (scipy.stats.ttest_ind) were applied to compare these metrics across models, with statistical significance differences determined using an alpha level of 0.05 adjusted by Bonferroni correction.

## Results

In this study, we investigated whether the discrepancy between DCNNs’ object recognition performance and neural predictability could be explained by scene segmentation processes. We trained 20 DCNNs to categorize objects under four distinct visual diets: original, object-only, background-silhouette, and background-only images (Figure 1). Through this systematic manipulation, we examined how varying scene segmentation demands affect both DCNNs’ categorization performance and alignment with neural responses.

### Object-only training enhances DCNNs’ categorization performance

First, we established that all models successfully learned object recognition, evidenced by converging training loss (Figure 2A) and above-chance validation performance (Figure 2B). Models trained on object-only images showed the most rapid learning and achieved the highest validation accuracy (∼53%), likely reflecting the reduced complexity when objects are presented in isolation without scene segmentation demands. Models trained on original and background-silhouette images achieved comparable intermediate performance (∼37% and ∼38%). The comparable performance between these conditions suggest that object shape information, combined with contextual background cues, are sufficient for effective categorization.

**Figure 2.**
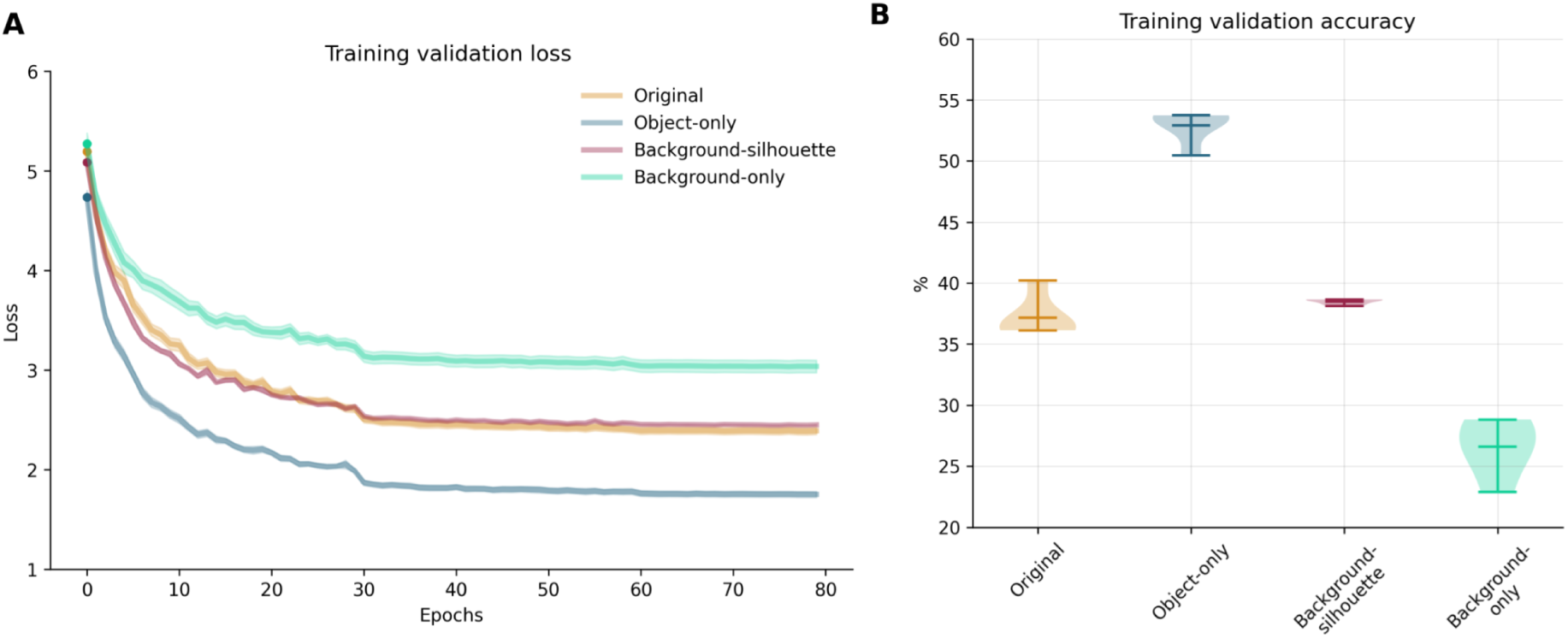
Training dynamics and categorization performance across visual diets. **A) Training validation loss across epochs for different visual diets.** Lines represent mean values across five model initializations per condition; shaded areas represent ±SEM. **B) Final validation accuracy for each image condition.** Violin plots reflect the median, 1st and 3rd quartiles across five model initializations; width of bars indicate density of values.

In contrast, background-only models showed the slowest learning and lowest accuracy (∼27%), though still performing above chance. This confirms that while contextual information alone carries meaningful signals for object categorization, the removal of explicit object information substantially impairs recognition performance, supporting theories about the importance of scene statistics in visual recognition (Oliva and Torralba, 2006, 2007) and also revealing its limitations. It is important to note that in the background-only condition, some residual shading from the original object shape occasionally remains (Figure 1D). As a result, the influence of these subtle cues on model performance cannot be entirely ruled out.

Table 1 details all statistical comparisons on models’ categorization performance between image conditions. All models significantly differ from each other except for models trained on original and background-silhouette images. These findings demonstrate a clear trade-off between scene complexity and object recognition performance. While removing contextual information (object-only condition) enables faster learning and higher accuracy, it may result in less robust representations that don’t capture the full complexity of natural vision. Conversely, the ability of models to learn from background information alone, albeit with reduced performance, suggests that contextual processing plays a more fundamental role in visual recognition than previously appreciated. This pattern of results sets the stage for examining whether these differences in categorization performance correspond to differences in neural prediction accuracy, potentially helping explain the observed disconnect between DCNNs’ categorization performance and neural alignment.

**Table 1.** Pairwise comparisons on models’ object categorization performance.

| Conditions compared | t-statistic | p-value |
| --- | --- | --- |
| Original vs Object-only | -15.97 | <0.001* |
| Original vs Background-silhouette | 1.69 | 0.129 |
| Original vs Background-only | 7.95 | <0.001* |
| Object-only vs Background-silhouette | 23.02 | <0.001* |
| Object-only vs Background-only | 21.07 | <0.001* |
| Background-silhouette vs Background-only | 10.90 | <0.001* |

\* indicates statistical significance based on a Bonferroni corrected alpha value of 0.05

### EEG encoding performance is uniformly high despite varying categorization performance

To investigate the relationship between categorization ability and neural prediction, we built encoding models using representations from our differently trained DCNNs to predict EEG responses. We included five ImageNet-trained and five randomly initialized AlexNets as control conditions, providing benchmarks for both task-optimized and untrained networks.

Remarkably, all models demonstrated uniformly high EEG encoding performance despite their substantial differences in categorization accuracy (27-53%, Figure 2B). The temporal dynamics of neural prediction revealed a characteristic peak at ∼0.1s post-stimulus onset (Figure 3A), with models trained on original and background-only images approaching the lower bound of the noise ceiling. This early peak timing suggests that all models represented fundamental visual features during initial scene analysis.

**Figure 3.**
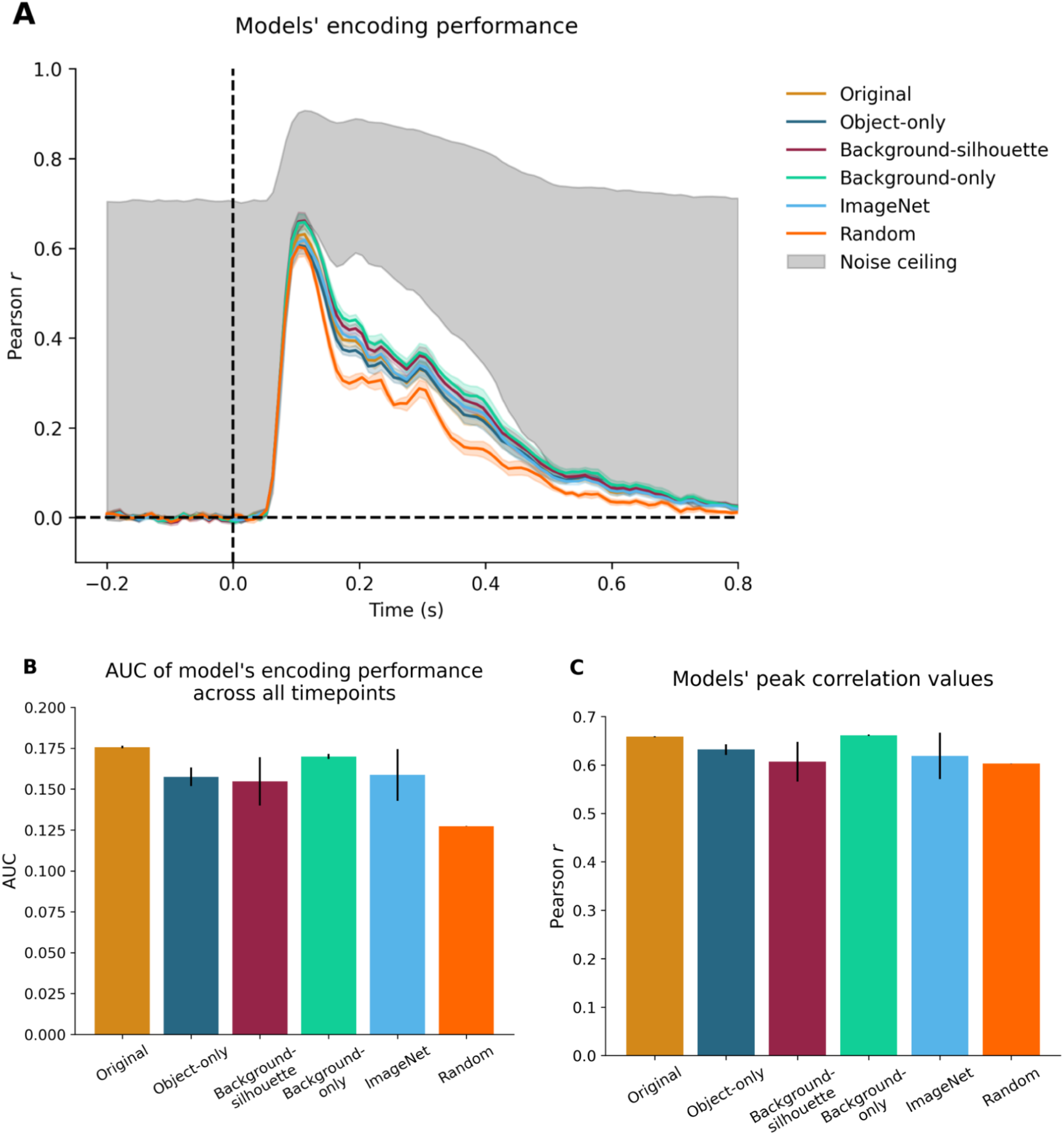
Model encoding performance across visual diets. **A) Temporal dynamics of neural prediction.** All models show peak encoding performance at ∼0.1s post-stimulus onset, followed by a decrease. Lines show mean correlation across 10 subjects; shaded areas represent standard error. **B) Area Under the Curve (AUC) of encoding performance across full time window (0-0.8s).** Bars represent mean AUC across subjects; error bars indicate standard error. **C) Peak correlation values of encoding performance.** Bars represent mean Pearson *r* across subjects; error bars indicate standard error.

Beyond this initial peak, we observed a systematic decline in encoding performance across all models between 0.15s and 0.5s. This decline was particularly pronounced in random/untrained models, which showed significantly greater divergence from trained models during later processing stages. These results suggest that task-specific training enhances alignment with EEG signals in later temporal windows, likely refining representations that support mid-to late-stage visual processing.

To quantify encoding performance, we examined both the Area Under the Curve (AUC) and peak correlation values across all conditions. The AUC analysis (Figure 3B) showed that while the original model achieved the highest value (0.176), followed closely by background-only (0.170) and object-only (0.158) models, these differences were not statistically significant. Original and background-only models significantly outperformed random models (*p* < 0.001, Bonferroni-corrected) but have comparable performance with all other trained models, providing evidence that models’ encoding performance benefits from training but is independent of object-specific learned features. Notably, models trained on ImageNet showed comparable performance, suggesting that our training on Google Open Images V5 dataset yielded representations which were equally effective at predicting EEG responses.

Peak correlation analyses (Figure 3C) showed that all trained models reached high peak values (ranging from 0.60 to 0.66). The background-only model reached the highest peak value (0.66) and the random model reached the lowest value (0.60). The small difference in peak correlations between the trained models and random models indicate that - 1) random models can capture early visual features to a large degree, and 2) training influences models’ encoding performance much more beyond the initial 0.1s peak. Additionally, the modest improvement of trained models over random initialized ones (0.06) suggests that while training enhances neural prediction, this enhancement is relatively subtle compared to the substantial differences in categorization performance. Original, background-only and object-only models significantly outperformed random models but also had comparable performance with all other trained models. This consistency further supports the hypothesis that early visual features drive EEG alignment.

These findings reveal a fundamental dissociation between object recognition capability and neural prediction accuracy. Despite differences in training and object categorization accuracy, DCNNs showed uniformly high encoding accuracy, suggesting that EEG signals predominantly reflect early visual processing features shared across visual diets, rather than high-level categorical representations. This pattern held true across multiple analysis approaches and temporal windows, providing robust evidence that neural responses captured by EEG primarily reflect fundamental scene processing mechanisms rather than learned categorical distinctions.

### Scene segmentation demands influence neural encoding

Our systematic manipulation of DCNN’s training visual diets revealed important patterns in how scene processing affects neural encoding. Models trained with complete scenes and therefore higher segmentation demands (original and background-only conditions) consistently outperform untrained models in neural prediction (*p* < 0.001, Bonferroni-corrected), even though this background-only condition showed the lowest object categorization accuracy (∼27%). This finding suggests that exposure to full scene complexity, which necessitates segmentation processes, enhances alignment with neural responses.

Interestingly, despite substantial differences in categorization accuracy between trained conditions (ranging from 27% to 53%), the differences in encoding performance between trained models did not reach statistical significance after correction for multiple comparisons (Table 2). This relatively uniformity in neural prediction across training conditions stands in striking contrast to their divergent categorization abilities.

**Table 2.** Pairwise comparisons of models’ encoding performance – AUC.

SCENE SEGMENTATION PROCESSES DRIVE EEG-DCNN ALIGNMENT
| Conditions compared | t-statistic | p-value |
| --- | --- | --- |
| Original vs Object-only | 3.17 | 0.01 |
| Original vs Background-silhouette | 1.42 | 0.19 |
| Original vs Background-only | 3.36 | 0.01 |
| Original vs ImageNet | 1.08 | 0.31 |
| Original vs Random | 59.75 | <0.001* |
| Object-only vs Background-silhouette | 0.18 | 0.86 |
| Object-only vs Background-only | -2.12 | 0.07 |
| Object-only vs ImageNet | -0.07 | 0.95 |
| Object-only vs Random | 5.30 | <0.001* |
| Background-silhouette vs Background-only | -1.03 | 0.33 |
| Background-silhouette vs ImageNet | -0.18 | 0.86 |
| Background-silhouette vs Random | 1.85 | 0.10 |
| Background-only vs ImageNet | 0.71 | 0.50 |
| Background-only vs Random | 28.30 | <0.001* |
| ImageNet vs Random | 1.98 | 0.08 |

**Table 3.** Pairwise comparisons of peak correlation values between models.

| Conditions compared | t-statistic | p-value |
| --- | --- | --- |
| Original vs Object-only | 2.45 | 0.04 |
| Original vs Background-silhouette | 1.25 | 0.25 |
| Original vs Background-only | -1.61 | 0.15 |
| Original vs ImageNet | 0.83 | 0.43 |
| Original vs Random | 65.76 | <0.001* |
| Object-only vs Background-silhouette | 0.58 | 0.58 |
| Object-only vs Background-only | -2.70 | 0.03 |
| Object-only vs ImageNet | 0.27 | 0.80 |
| Object-only vs Random | 2.66 | 0.03 |
| Background-silhouette vs Background-only | -1.32 | 0.22 |
| Background-silhouette vs ImageNet | -0.19 | 0.86 |
| Background-silhouette vs Random | 0.10 | 0.92 |
| Background-only vs ImageNet | 0.89 | 0.40 |
| Background-only vs Random | 34.52 | <0.001* |
| ImageNet vs Random | 0.33 | 0.75 |

\*\*\* indicates statistical significance based on a Bonferroni corrected alpha value of .05

These findings suggest that the fundamental visual features driving EEG signals may be more basic and universal than the specialized features enabling successful object recognition. Rather than a linear relationship between segmentation demands and neural encoding, we observed a more nuanced pattern where all models appear to capture many essential visual processes reflected in EEG responses, with subtler enhancements from complete scene processing.

### Layer-wise analysis reveals dissociation between categorization and neural encoding

To directly test whether scene segmentation or object categorization processes drive EEG responses, we analyzed how different DCNN layers contribute to both functions. If encoding performance primarily reflects object categorization processes, we would expect deeper layers (which better distinguish object categories) to show stronger EEG prediction. Conversely, if encoding performance reflects scene segmentation processes, earlier layers (which extract and segregate low- and mid-level visual features) should better predict EEG responses.

We performed a layer-wise decoding of object categories on our fine-tuned models and plotted each layer’s decoding accuracy against its peak correlation with EEG responses (Figure 4). Note that the layer-wise decoding is performed on the fine-tuned models because these are the models used to predict EEG responses. This approach revealed three clear patterns: First, the models’ ability to distinguish between object categories generally improved from shallow to deeper network layers, with performance peaking at the maxpool5 layer (Figure 4A). This progression was consistent across all trained models, though with varying magnitudes - models trained on complete scenes (original condition) maintained more robust representations in deeper layers compared to other conditions, suggesting the importance of contextual information for stable object representations.

**Figure 4.**
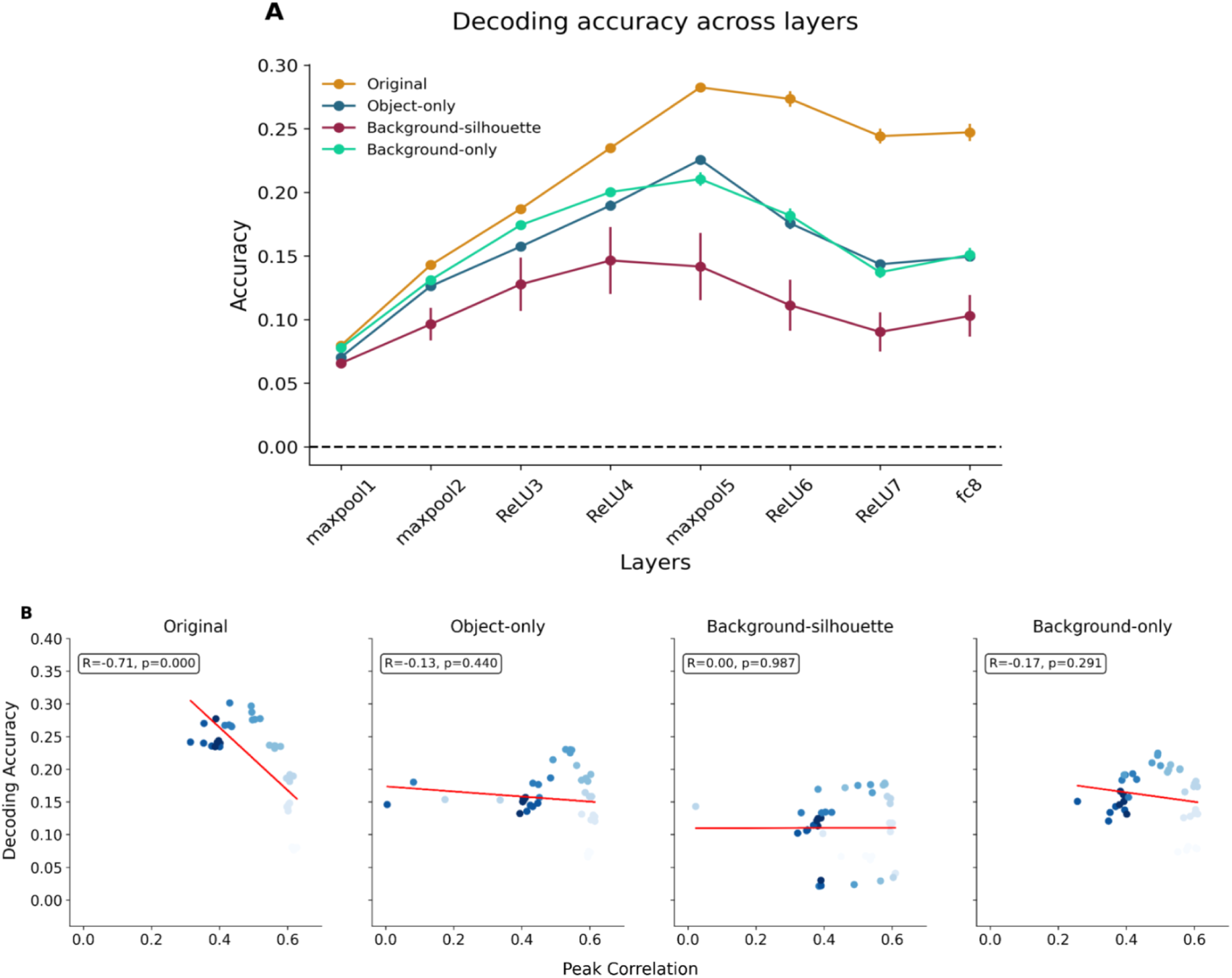
Layer-specific feature analysis and EEG encoding performance. **A) Object category decoding accuracy across network layers for different training conditions.** Fine-tuned DCNNs trained under different visual diets exhibit different abilities to distinguish between object categories, with performance peaking at maxpool5. Lines represent mean accuracy across five model initializations; error bars indicate standard error. **B) Relationship between layer-wise categorization accuracy and encoding performance.** Scatter plots show decoding accuracy versus peak EEG encoding correlation for each layer. Point color indicates layer depth (lighter = earlier layers). Negative correlations indicate dissociation between categorization ability and encoding performance.

Second, we observed striking negative correlations between layers’ categorization accuracy and their encoding performance. This inverse relationship was strongest in models trained on original images (*r* = -0.71, *p* < 0.001), with object-only, background-silhouette and background-only models showing weaker but still negative correlations. The color gradient visualization of layer depth in our scatter plots (see Figure 4B) shows earlier layers (lighter-colored points) clustering toward higher encoding performance, while deeper layers (darker points) show higher categorization accuracy but lower encoding performance.

Third, this dissociation proved robust across all training conditions, with earlier layers consistently showing better prediction of neural responses despite their limited categorical knowledge. This pattern strongly supports our hypothesis that encoding models predominantly capture EEG responses related to fundamental visual processing rather than categorical representations.

Together, these findings strongly support our hypothesis that encoding models of EEG responses predominantly capture scene segmentation processes whether the visual system extracts and organizes basic image properties involving early and mid-level visual features, rather than high-level categorical representations.

## Discussion

### Dissociation between object recognition and neural encoding

Our results confirmed and extended previous findings that DCNNs’ object categorization performance can be decoupled from their encoding performance of neural data (Xu and Vaziri-Pashkam, 2021; Linsley et al., 2023). Despite varying object categorization accuracy (27-53% for Top-1 accuracy), all trained models showed remarkably similar EEG encoding performance. Most notably, our layer-wise analysis revealed a significant negative relationship between categorization accuracy and encoding performance, particularly in models trained on original images (*r* = -0.71, *p* < 0.001). This finding challenges the traditional assumption that better object categorization performance necessarily leads to better neural prediction. Instead, it provides strong quantitative evidence that features distinguishing between object categories do not drive a large portion of EEG responses. This dissociation between object categorization and neural encoding highlights the importance of explicitly evaluating encoding models.

The consistent peak in encoding performance at ∼0.1s post-stimulus onset across all models, including random models, suggests that DCNNs’ representations capture early visual processing stages. This timing aligns with previous research showing rapid processing of low and mid-level features in the visual system (VanRullen and Thorpe, 2001; Scholte et al., 2009). Our findings caution against over-interpretation of EEG decoding studies. The successful decoding of object categories from EEG likely reflects the correlation between low and mid-level visual features and semantic categories rather than direct access to high-level categorical representations.

### Scene segmentation as a fundamental process

Our systematic manipulation of visual diets revealed the role of scene segmentation in driving neural responses, helping explain the disconnect between DCNNs’ categorization performance and neural prediction. Despite achieving the lowest categorization accuracy, models trained on background-only images demonstrated encoding performance comparable to those trained on original images, with both conditions showing peak correlations around 0.1s post-stimulus. Models trained on complete scenes (original and background-only conditions) - which have higher segmentation demands - demonstrated better encoding performance compared to those on simplified inputs (object-only and background-silhouette conditions). This pattern suggests that exposure to the full complexity of natural environments during training promotes the development of representations that better align with biological visual processing. These findings align with evidence from human vision, where rapid scene understanding relies heavily on global image properties and contextual cues (Oliva and Torralba, 2006; Greene and Oliva, 2009; Groen et al., 2017) and highlight the significant role of scene statistics and contextual features in driving early neural responses, even in the absence of explicit object information. The superior performance of models trained with complete scenes support theories emphasizing the importance of scene segmentation and recurrent processing (Scholte et al., 2008) and aligns with evidence that object recognition in natural environments relies heavily on contextual processing (Peelen and Kastner, 2011). Our findings highlight an important principle: effective visual processing requires not just the ability to recognize isolated objects, but also the capacity to interpret them within their natural contexts. The fact that models trained with higher segmentation demands showed more generalizable performance (see Figure 4A) suggests that learning to process complete scenes leads to more robust visual representations, even if this comes at the cost of lower initial categorization accuracy.

### Theoretical Implications for Modeling Visual Processing

Our results contribute to the ongoing discussion about how DCNNs relate to biological vision, suggesting this relationship may be more complex than previously thought. While DCNNs are primarily feedforward architectures, the visual features they learn appear to capture aspects of both feedforward and recurrent processing in biological vision. The brain’s visual system is highly recurrent, with feedback connections playing crucial roles in scene segmentation, figure-ground segregation, and object recognition (Lamme and Roelfsema, 2000; Self et al., 2013). Scene segmentation, in particular, is known to rely heavily on recurrent processing in biological vision, as the brain must iteratively refine its interpretation of object boundaries and relationships within complex scenes (Scholte et al., 2008; Seijdel et al., 2020).

The strong encoding performance we observe at early time points (∼0.1s) likely reflects both initial feedforward processing and the early stages of recurrent computations that are essential for scene segmentation; this finding is congruent with outcomes of previous experiments (Loke et al., 2022). Our finding that models trained with higher segmentation demands better predict neural responses is particularly intriguing, as these models may be developing representations that capture computations that typically require recurrent processing in biological systems.

The superior encoding performance of earlier network layers takes on new significance in this context. While these layers are often thought to represent simple feedforward features, their ability to predict neural responses may stem from capturing computational elements that arise from both feedforward and recurrent processing in biological vision. This interpretation aligns with recent work showing that feedforward DCNNs can sometimes approximate computations that require recurrence in biological systems (Kar et al., 2019; Kietzmann et al., 2019), though they likely do so by compressing temporal dynamics into spatial hierarchies.

This could have important implications for interpreting EEG decoding studies: successful category decoding may reflect not just feedforward processing of low-level features, but also early recurrent computations involved in scene segmentation that begin to emerge around 100ms post-stimulus onset. Our findings suggest that even strictly feedforward DCNNs can capture aspects of recurrent processing in biological vision, particularly those involved in scene segmentation, though they likely achieve this through fundamentally different computational mechanisms. Understanding these relationships between artificial and biological visual processing, especially regarding how feedforward networks can approximate recurrent computations, remains a crucial challenge for future research

### Limitations and future directions

Our findings need to be interpreted under several limitations. First, while AlexNet provided computational efficiency and established performance benchmarks, future work should examine whether our findings generalize across different DCNN architectures, particularly more modern ones incorporating attention mechanisms or recursive processing. Second, while EEG provides high temporal resolution, it provides limited spatial information about visual processing and only captures summated neural activity. Third, our background-only condition, while innovative, may retain subtle shading cues from removed objects. Future work could employ more sophisticated image manipulation techniques to more completely isolate contextual effects. Finally, the usage of rapid serial visual presentation in the experimental paradigm for EEG recordings may mask some aspects of neural processing. Future research should explore how these findings generalize across different DCNN architectures, neuroimaging modalities and experimental paradigms.

## Conclusion

Our study provides new insights into why improvements in DCNNs’ object recognition performance don’t necessarily translate to better neural prediction. Through systematic manipulation of scene segmentation demands in DCNNs, we demonstrated that EEG signals predominantly reflect fundamental scene parsing processes as captured in early DCNNs processing layers rather than high-level object representations. This finding was supported by two key observations: first, the uniformity in EEG encoding performance despite substantial variation in categorization accuracy (27-53%), and second, the significant negative correlation between layer-wise categorization ability and encoding performance.

Our results reveal that the ability to predict neural responses depends more on capturing fundamental scene segmentation processes than on object recognition capability. This helps us explain why modern DCNNs, despite their impressive categorization performance, may not show corresponding improvements in neural prediction - they may be optimizing features that diverge from basic scene parsing mechanisms. The consistent peak in encoding performance at ∼0.1s post-stimulus onset across all models, including those trained only on backgrounds, suggests that these early neural responses primarily reflect the initial stages of scene segmentation, where the visual system extracts and organizes scene information. Rather than being merely a prerequisite for object recognition, scene segmentation could be a more fundamental aspect of visual processing than previously recognized. Future work should investigate how these findings generalize across different neural measurement modalities.

